# Whole-body imaging of neural and muscle activity during behavior in *Hydra*: bidirectional effects of osmolarity on contraction bursts

**DOI:** 10.1101/2019.12.20.883835

**Authors:** Wataru Yamamoto, Rafael Yuste

## Abstract

The neural code relates the activity of the nervous system to the activity of the muscles to the generation of behavior. To decipher it, it would be ideal to comprehensively measure the activity of the entire nervous system and musculature in a behaving animal. As a step in this direction, we used the cnidarian *Hydra vulgaris* to explore how physiological and environmental conditions alter the activity of the entire neural and muscle tissue and affect behavior. We used whole-body calcium imaging of neurons and muscle cells and studied the effect of temperature, media osmolarity, nutritional state and body size on body contractions.

In mounted *Hydra*, changes in temperature, nutrition or body size did not have a major effect on neural or muscle activity, or on behavior. But changes in media osmolarity altered body contractions, increasing them in hipo-osmolar media solutions and decreasing them in hyperosmolar media. Similar effects were seen in ectodermal, but not in endodermal muscle. Osmolarity also bidirectionally changed the activity of contraction bursts neurons, but not of rhythmic potential neurons.

These findings show osmolarity-dependent changes in neuronal activity, muscle activity, and contractions, consistent with the hypothesis that contraction burst neurons respond to media osmolarity, activating ectodermal muscle to generate contraction bursts. This dedicated circuit could serve as an excretory system to prevent osmotic injury. This work demonstrates the feasibility of studying the entire neuronal and muscle activity of behaving animals.

**Significance Statement:** We imaged whole-body muscle and neuronal activity in *Hydra* in response to different physiological and environmental conditions. Osmolarity bidirectionally altered *Hydra* contractile behavior. These changes were accompanied by corresponding changes in the activity of one neuronal circuit and one set of muscles. This work is a step toward comprehensive deciphering of the mechanisms of animal behavior by measuring the activity of all neurons and muscle cells.

## Introduction

The introduction of calcium imaging of neuronal circuits (Yuste and Katz 1991) has enabled recent investigations of the circuit basis of animal behavior in a number of transparent organisms such as *C. elegans, Drosophila* larvae and zebrafish embryos (Nagel, Brauner et al. 2005, Liewald, Brauner et al. 2008, Honjo, Hwang et al. 2012, Cong, Wang et al. 2017, Kim, Kim et al. 2017). While these studies have focus on a particular part of the nervous system, in order to systematically understand the neural code, i.e., the relation between the activity of a nervous system and behavior, it would be ideal to measure the activity of the entire nervous system and the entire muscular tissue during the entire behavioral repertoire of an animal. This could be possible with the transparent fresh-water cnidarian *Hydra vulgaris*, using transgenic strains that express calcium indicators in every neuron (Dupre and Yuste 2017) and every muscle cell of the body (Szymanski and Yuste 2019), and applying machine learning to systematically analyze its behavior (Han, Taralova et al. 2018). *Hydra* has one of the simplest nervous system in evolution, with several hundreds to a few thousand neurons (Hadzi 1909, Parker 1919, Westfall, Wilson et al. 1991), organized in two independent nerve nets, in the ectoderm and endoderm (Dupre and Yuste 2017). *Hydra*’s nerve nets have sensory cells and ganglion cells and are distributed throughout the body of the animal, without any cephalization or ganglia (Epp and Tardent 1978). This simplicity gives hope that a systematic measurements of the entire neural and muscular activity of behaving *Hydra* could be used to decipher systematically the mechanisms of behavior.

As a step in this direction, we focused on a simple contractile behavior, and used calcium imaging to measure how neurons and muscular cells in mounted and freely-behaving *Hydra* responds to physiological and environmental conditions that are important for their survival. ^Experimental conditions included high or low osmolarity (50mM sucrose or diH_2_O), temperature^ (23°C or 30°C), amount of food (from 0 to 4 shrimp/day), and body size (mature vs. newly released buds). We find that motor behaviors such as contractions and locomotion are readily altered by osmolarity in a bidirectional manner. Corresponding changes are found in the activity of ectoderm muscle and contraction burst neurons. Other conditions did not alter the activity of ectoderm muscle and neurons. Our findings indicate that *Hydra’s* contraction burst circuit senses osmolarity to control ectodermal muscle and generate contractile behaviors.

## Material and Methods

### Materials

Sucrose and Sea Salt were purchased from Sigma. Brynn shrimp, *Artemia nauplii* was obtained from Brine Shrimp Direct. We used transgenic *Hydra* expressing GCaMP6s in neurons (Dupre and Yuste 2017) or ectoderm/endoderm muscle cells (Szymanski and Yuste 2019).

### Hydra *culture*

*Hydra* were maintained in media composed of 1.3mM CaCl_2_, 0.02mM MgCl_2_, 0.03mM KNO_3_, 0.5mM NaHCO_3_, 0.08mM MgSO_4_ in an 18°C incubator. *Hydra* were fed with brine shrimp three times a week and were starved for two days prior to an experiment.

### Environmental or Physiological Conditions

The following conditions were used:

(1) Food: *Hydra* were fed 0, 1 or 4 shrimps every day for a week.

(2) Size: *Hydra* with a large body size (~1 cm) or a small size body (~0.3 mm, fed once after the bud separated from its adult *Hydra*) were chosen.

(3) Temperature: room (23°C) or high temperature (30°C).

(4) Osmolarity: *Hydra* were imaged in media with low (diH_2_O, 0 mOsm/L), medium (control, *Hydra* media, 5mOsm/L, fresh water is usually between 2-8 mOsm/L), or high (50mM Sucrose, 50mOsm/L) osmolarity.

### Calcium imaging

Wide-field calcium imaging of *Hydra* was conducted at 2 Hz using a fluorescence dissecting microscope (Leica M165) equipped with a long-pass GFP filter set (Leica filter set ET GFP M205FA/M165FC), 1.63X Plan Apo objective, and a sCMOS camera (Hamamatsu ORCA-Flash 4.0). A mercury arc lamp was used to illuminate the sample. *Hydra* were mounted between coverslips with 100-200 um spacer, depending on its thickness. All imaging were conducted at a room temperature ~23°C unless indicated.

### Behavior test

The number of contractions and foot detachment were manually scored from calcium imaging movies used (mounted *Hydra* between coverslips) or movies of freely moving *Hydra* in glass-bottom dishes (MatTek). Five animals were placed per well (depth is 700-750 um) for 1 hour recordings.

### Analysis of neural and muscular activity

Values for whole-body fluorescent intensity in each frame over time were obtained with ImageJ and used to detect peaks of the CB and RP1 pulses using a semi-automated program in MATLAB. Whole-body muscle activity was analyzed in the same manner.

### Measuring width of body column

*Hydra* were imaged at 0.5 Hz using a dissecting microscope (Leica M165), 1.63X Plan Apo objective, and sCMOS camera (Hamamatsu ORCA-Flash 4.0). *Hydra* were mounted between coverslips with around 200um spacer in control media, or in high osmolarity solution (50mM Sucrose). To measure width, the body column of *Hydra* was fitted into ellipse using a program written by MATLAB. The lowest values from each cycle were taken to calculate average width at the end of the elongation.

### Statistical Methods

Data are shown as average ± SEM in figures and in the text. Two-tailed unpaired student t test or one-way ANOVA with Tukey multiple comparison test were conducted in GraphPad Prism software.

### Code Accessibility

The code will be made available.

## Results

### *Hydra’s* contractile behavior affected by media osmolarity

*Hydra* has a small repertoire of highly stereotypical behaviors (Han, Taralova et al. 2018). One of the most noticeable ones are spontaneous periodic contractions, known as “contraction bursts” (Wagner 1905, Reis and Pierro 1944, Passano and Mccullough 1964). Possible roles of contractions by *Hydra* include foraging and protection by retraction (Miglietta, Della Tommasa et al. 2008, Swain, Schellinger et al. 2015), food digestion (Shimizu and Fujisawa 2003), and removing excess water from the body (Macklin, Roma et al. 1973). Another common behavior of *Hydra* is locomotion, i.e. translocation of the foot from one place to another. This is initiated by ‘foot detachment’ where the basal disk detaches from a substrate’s surface (Rodrigues, Ostermann et al. 2016). Here we tested how these two common behaviors of *Hydra* are affected by various physiological and environmental conditions. Conditions chosen included amount of food, osmolarity or temperature of media, and the size of an animal. For each condition, the frequency and duration of contractions and foot detachments were measured.

In mounted preparations, were specimens are place in a microscope chamber with a spacer, osmolarity or body size changed the frequency of contractions (Fig. 1A, B and C; see Methods). High osmolarity media significantly decreased the frequency of contractions compared to control (Fig. 1B, p = 0.0380) or low osmolarity (Fig. 1B, p = 0.0367). Similarly, high osmolarity media significantly decreased the number of foot detachment compared to control (Fig. 1C, p = 0.0003) or low osmolarity (Fig. 1C, p < 0.0001). Also, smaller size *Hydra* had more contraction (Fig. 1B, p = 0.0008) and fewer foot detachment (Fig. 1C, p = 0.0378).

**Figure 1.**
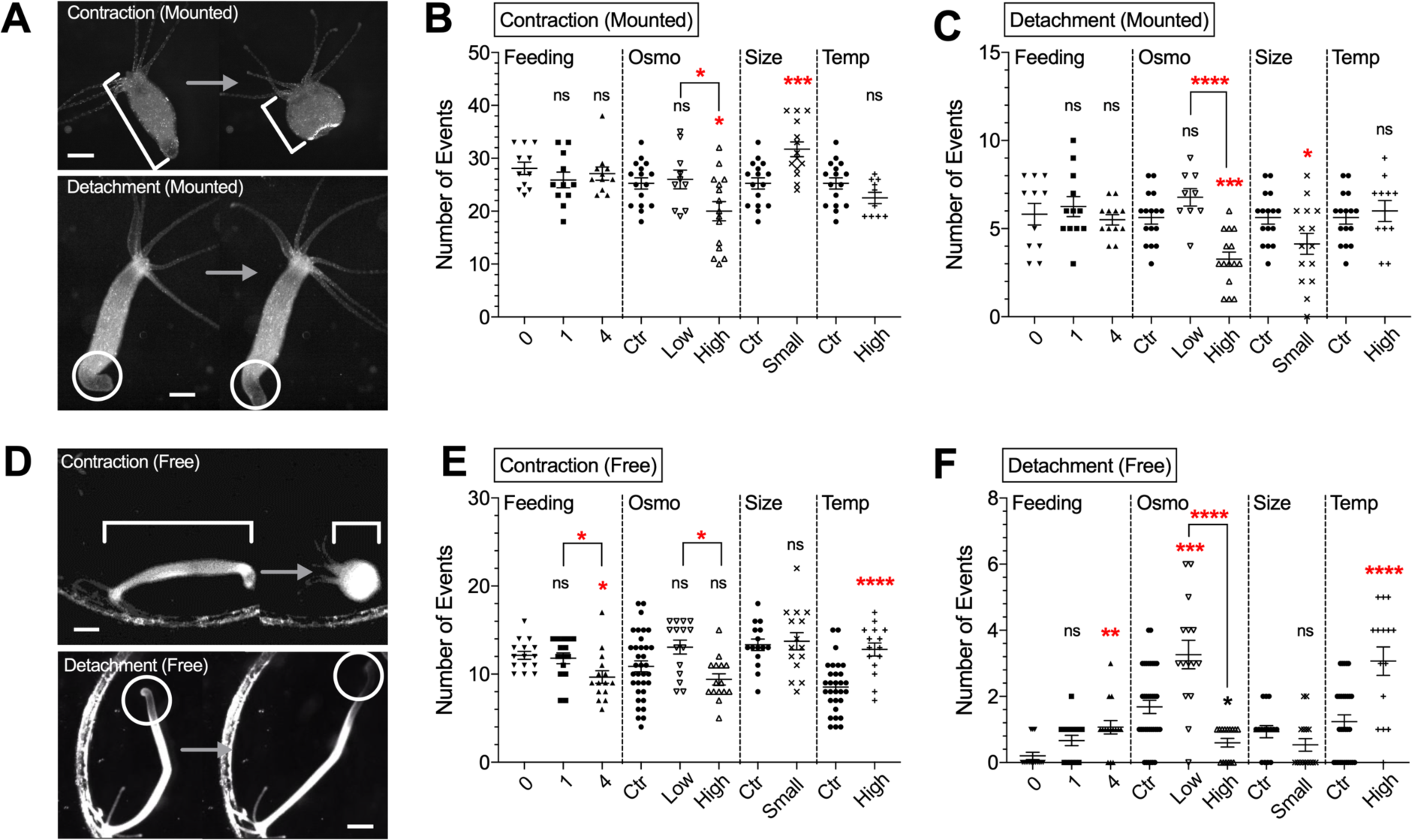
Contraction and locomotion behavior under different experimental conditions. Movies from mounted preparations during calcium imaging used for Figure 1 and Figure 2 analyzed in ***A***–***C***, and movies from 1-hour free-moving *Hydra* analyzed in ***D***–***F***. ***A***, Upper images depict changes in body length during longitudinal contraction. Lower images depict foot detachment. Scale bar, 500 µm. Number of contraction (***B***) and foot detachment (***C***) were counted. ***D***, Upper images depict changes in body length during longitudinal contraction. Lower images depict foot detachment followed by locomotion. Scale bar, 1 mm. Number of contraction (***E***) and foot detachment/locomotion (***F***) were counted. Abbreviations: Osmo, osmolarity; Temp, Temperature. Error bars shown as mean ± SEM, with symbol marks denoting data points from individual *Hydra* (*N* = 9–16 for ***B*** and ***C***; *N* = 15–30 for ***E*** and ***F***). Tukey multiple comparisons tests were performed following one-way ANOVA for osmolarity experiment, and Student t test was performed for others: ns ≥ 0.05, \**p* < 0.05; \*\**p* < 0.01; \*\*\**p* < 0.001; \*\*\*\**p* < 0.0001.

As mounting restricts *Hydra* behavior, due to compression of body between glass coverslips, we also imaged freely moving *Hydra* under widefield illumination in the same conditions (Movie 1). Consistent with results in mounted preparations (Figure. 1B and C), in free moving animals, high osmolarity also decreased the number of contractions compared to low osmolarity (Fig. 1E, p = 0.0100) and the number of foot detachments, compared to control (Fig. 3F, p = 0.0134) or low osmolarity (Fig. 3F, p < 0.0001). Unlike mounted preparations, in freely moving animals, feeding (4 shrimps per day) decreased the number of contraction (Fig. 1E, p = 0.0164) while increasing foot detachments (Fig. 1F, p = 0.0014). High temperature also increased contractions (Fig. 1E, p < 0.0001) and foot detachments (Fig. 3F, p < 0.0001). Overall, osmolarity was the only parameter that robustly changed behavior in both freely moving and mounted specimens. As motor behaviors must be generated as a result of contractile force derived from muscle, we next assessed how these changes in behaviors are accounted for the activity of muscle cells. For these experiments, we used exclusively mounted preparation, as it is yet not feasible to image and reconstruct the activity of neurons and muscle cells in freely moving animals.

### Bidirectional effects of osmolarity on ectodermal muscle activity

*Hydra*’s body is composed of two layers of cells: ectodermal and endodermal epitheliomuscular tissues. Both epithelia are separated by an extracellular matrix called mesoglea. Inside these epithelial layers there is a gastrovascular cavity that functions as a gut and vasculature that carry nutrients to the entire body (Shimizu and Fujisawa 2003). Both ectoderm and endoderm muscle generate action potentials (Dupre and Yuste 2017, Szymanski and Yuste 2019), which likely propagate through gap junctions (Westfall, Kinnamon et al. 1980). These muscle cells generate motions in a calcium-dependent manner through myonemes, intracellular muscle processes that run longitudinally in the ectoderm and radially in the endoderm (Otto 1977). Thus, *Hydra* generates motor behavior such as contractions and elongations by coordinating the activity of these two layers of muscle (Szymanski and Yuste 2019). However, how their activity is affected by physiological and environmental conditions has not been characterized. To test the effect of environmental manipulations on muscle activity, we used transgenic *Hydra* that express genetically-encoded calcium indicator GCaMP6s in every ectoderm or endoderm muscle cell (Szymanski and Yuste 2019). With these transgenic animals, 2-hour long calcium imaging sessions was conducted (Movie 2) to explore how each physiological or environmental condition changes muscle activity (Fig. 2A).

**Figure 2.**
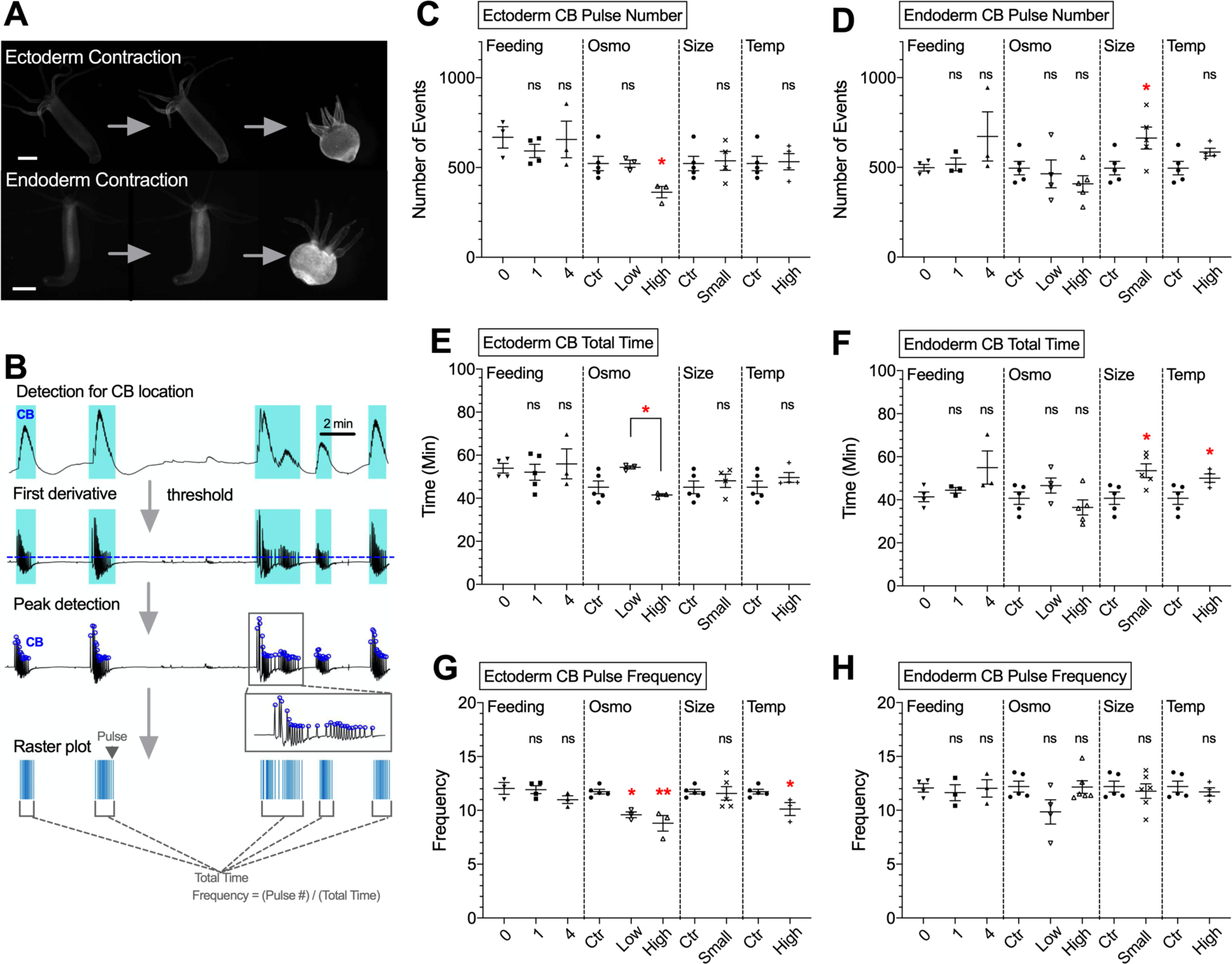
Analysis of ectoderm and endoderm muscle activity in under different experimental conditions. ***A***, Upper images depict contraction burst (CB) in *Hydra* expressing GCaMP6s in ectoderm muscle. Lower images depict CB in *Hydra* expressing GCaMP6s in endoderm muscle. Scale bar, 500 µm. ***B***, Schematic summarizing the steps to detect peaks of CB pulses from raw traces extracted from of 2-hour calcium imaging movies. RP1 pulses were not present in muscular activity. ***C***–***H***, Each parameter was analyzed using under four different conditions noted: ***C***, ectoderm CB pulse number; ***D***, endoderm CB pulse number; ***E***, ectoderm CB total time; ***F***, endoderm CB total time: ***G***, ectoderm CB total time; ***H***, endoderm CB total time. Abbreviations: Osmo, osmolarity; Temp, Temperature. Error bars are shown as the mean ± SEM, with symbol marks denoting data points from individual *Hydra* (*N* = 3–6). Tukey multiple comparisons tests were performed following one-way ANOVA for Osmolarity experiment, and Student t test was performed for others: ns ≥ 0.05, \**p* < 0.05.

Widespread activation of the entire body musculature was observed when *Hydra* contracts, as described (Szymanski and Yuste 2019), with transient calcium increases that synchronously occurs in the entire muscle tissue. These activations usually appeared as a burst during each contraction event, reflecting behavioral contraction bursts (CBs) (Passano and McCullough 1963, Passano and Mccullough 1964). To analyze the spatiotemporal dynamics of these muscle pulses and bursts, we used a computer program to semi-automatically detect events from whole body fluorescence intensity measurements (Fig. 2B). In agreement with behavioral data (Fig. 1), in ectoderm muscle tissue high osmolarity decreased the number of pulses (Fig. 2C, p = 0.0356), burst duration (Fig. 2E, p = 0.0273) and frequency (Fig. 2G, p = 0.0017), as compared to low osmolarity. In contrast, there was no change in endoderm muscle activity in response to osmolarity changes, although increases in endoderm muscle activity were observed in smaller *Hydra*, or with increased temperature (Fig. 2D, F and H).

We concluded that osmolarity changed ectodermal muscle activity in the same way as it changed contractile behavior, but did not affect endodermal muscle. We then examined the neural activity, supposedly upstream of this muscle activation.

### Bidirectional effect of osmolarity on contraction burst circuit activity

*Hydra* ‘s nerve nets lie at the base of both ectodermal and endodermal epithelial layers (Sarras, Meador et al. 1991) and is divided functionally into non overlapping circuits (Dupre and Yuste 2017). Two of such circuits are called contraction burst (CB) neurons and rhythmic potential 1 (RP1) networks (Dupre and Yuste 2017). These circuits activate in synchronous and oscillatory manner during *Hydra*’s spontaneous contraction (CB) or during elongation (RP1) (Passano and McCullough 1963, Rushforth and Burke 1971, Dupre and Yuste 2017). However, while these circuits likely have a combination of sensory and motor neurons, the exact role of these cells is still unclear. Similar to bilaterian species, the cnidarian *Hydra* has neuromuscular junctions (Chapman, Kirkness et al. 2010), and there is evidence suggesting direct interaction of muscle cells and neurons. First, gap junctions are found between muscle cells and neurons (Westfall, Kinnamon et al. 1980). Also, *Hydra* contractions are greatly reduced after chemically eliminating neurons (Campbell, Josephson et al. 1976), suggesting that muscle activity in *Hydra* are initiated and coordinated by neurons. We therefore set out to study neural activity in *Hydra* to account for the observed changes in the muscle activity and behavior under different conditions.

Similar to muscle imaging experiments (Fig. 2), 2-hour calcium imaging sessions were conducted using *Hydra* expressing GCaMP6s in the entire nerve net (Movie 3, Figure 3A) (Dupre and Yuste 2017). Then, the spatiotemporal dynamics of the CB and RP1 pulses were semi-automatically extracted using a computer program from whole-body fluorescence measurements (Fig. 3B), and events frequencies were calculated. Results showed that low osmolarity increased the number of CB pulses compared to control, while high osmolarity decreased them (p = 0.0422) compared to control or low osmolarity (p = 0.0005) (Fig. 3C), with no significant change in CB burst duration (Fig. 3E). Accordingly, high osmolarity decreased CB pulse frequency, compared to low osmolarity (p < 0.0001), while low osmolarity increased CB pulse frequency compared to controls (p = 0.0066) (Fig. 3G). Oher experimental conditions (feeding, temperature and body size) did not significantly alter the activity of CB neurons. These results indicate that CB neural activity is inversely proportional to osmolarity: lower osmolarity increases CB frequency while higher osmolarity decreases it.

**Figure 3.**
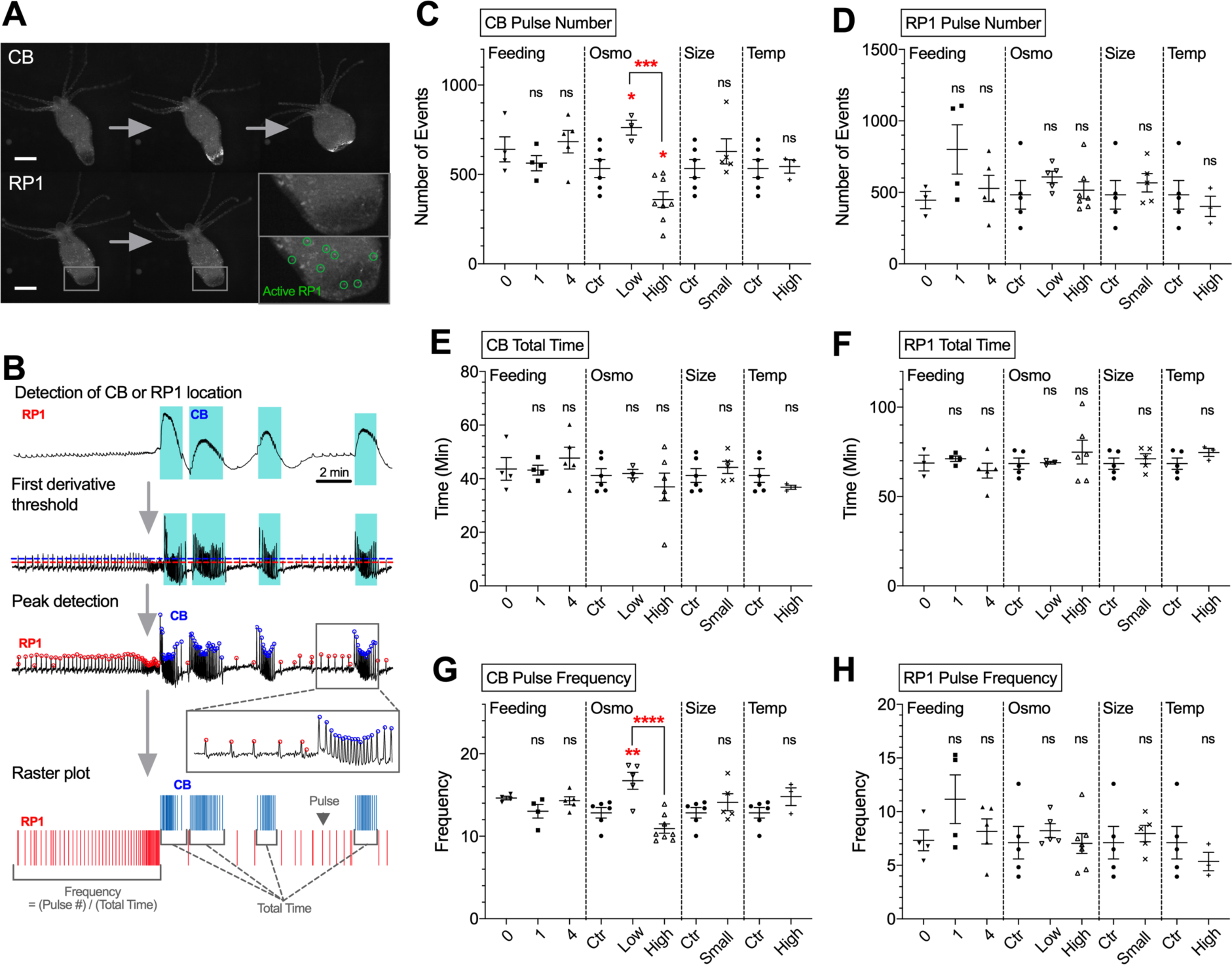
Analysis of CB and RP1 neuronal activity under different experimental conditions. ***A***, Upper images depict activation of contraction burst (CB) neurons. Lower images depict activation of rhythmic potential 1 (RP1) neurons. Scale bar, 500 µm. ***B***, Schematic summarizing steps to detect peaks of CB and RP1 pulses from raw traces extracted from 2-hour calcium imaging. ***C***–***H***, Each parameter was analyzed using under four conditions noted. Parameters analyzed were: ***C***, CB pulse number; ***D***, RP1 pulse number; ***E***, CB total time; ***F***, RP1 total time; ***G***, CB pulse frequency; ***H***, RP1 pulse frequency. Abbreviations used are: Osmo, osmolarity; Temp, Temperature. Error bars are shown as the mean ± SEM, with symbol marks denoting data points from individual *Hydra* (*N* = 3–8). Tukey multiple comparisons tests were performed following one-way ANOVA for Osmolarity experiment, and Student t test was performed for others: ns ≥ 0.05, \**p* < 0.05; \*\**p* < 0.01; \*\*\**p* < 0.001; \*\*\*\**p* < 0.0001.

In contrast to these results in CB neurons, none of the condition altered the activity of RP1 neurons, thought to be responsible for body elongation (Fig. 3D, F and H) (Dupre and Yuste 2017). These results suggest that the activity of RP1 neurons are apparently not affected by the environmental conditions tested. Overall, osmolarity consistently altered contractions, ectoderm muscle activity and CB neuronal activity, with hipo-osmolarity leading to increases and hyperosmolarity to decreases in all these three physiological outputs.

## Discussion

In this study, we examined the effect of internal and external experimental factors on the motor behavior and activity of muscle and neural tissue of *Hydra vulgaris*. We established imaging and analysis methods to measure the activity of neuron and muscle cells during behavior in mounted preparations, under different physiological and environmental conditions. Among the conditions tested (amount of food, osmolarity or temperature of media, and size of animal), osmolarity consistently affected three functional readouts: contractile behavior, ectoderm muscle activity and neural activity of the CB circuit. For foot detachments, ectodermal muscle CB duration and neuronal CB frequency, these effects were bidirectional, inversely related to osmolarity. Thus, *Hydra* can respond to osmolarity by specifically changing its neural and muscular activity, which presumably then changes behavior.

In both mounted and freely moving preparations, the number of contractions of *Hydra* in high osmolarity decreased compared to low osmolarity (Fig. 1B and E). This is consistent with previous findings that *Hydra’s* contraction rate is inversely proportional to media osmolarity (Benos and Prusch 1973). Changes of *Hydra* behavior with osmolarity are thought to be triggered by increased water accumulation in *Hydra’s* gastrovascular cavity, causing *Hydra* to swell. As *Hydra* cells are highly permeable to water (Lilly 1955), water could follow the concentration gradient between media (~5mOsm/L) and *Hydra* tissue (~120 mOsm/L), accumulating in the gastrovascular cavity (~60 mOsm/L), which serves as an excretory pathway in these basal metazoans that lack excretory systems (Benos and Prusch 1972). Furthermore, previous reports have suggested that the speed of water accumulation in *Hydra* depends on osmolarity (Kucken, Soriano et al. 2008, Soriano, Rudiger et al. 2009). Using regenerating hollow spheres of *Hydra* tissue fragments, made of two epithelial layers as in intact *Hydra*, the speed of sphere swelling due to water accumulation decreased linearly with increasing osmolarity (Kucken, Soriano et al. 2008, Soriano, Rudiger et al. 2009).

What are the mechanisms by which *Hydra* alters the contractions with osmolarity? One possibility is a mechanosensory system that could sense tissue pressure. Mechanosensory responses in *Hydra* have been characterized in cnidocytes (Kass-Simon and Scappaticci 2002), which use neurons to regulate their activation. *Hydra* is expected to express a set of potential osmoregulatory genes and mechanosenseory receptor genes, and it will be interesting to examine the functions of these proteins in regulating neuronal and muscular activity during behavior.

We propose the following model (Fig. 4A): *Hydra* undergoes a spontaneous cycle of elongation and contraction. In low osmolarity, this cycle speeds up due to increases in water accumulation and activation of mechanosensory receptors in the tissue. In contrast, in high osmolarity, this cycle slows down due to decrease in water accumulation and lesser activation of mechanosensory receptors. Consistent with this model, high osmolarity solution (50mM sucrose) significantly shortens the width of the body column, as if water accumulation were reduced (Fig. 4B-D). According to our results, body contractions would be generated by ectodermal muscles, themselves under the control of CB neurons. But while responses were indeed altered in an osmolarity-dependent manner in both CB neurons and ectoderm muscle tissue, our data also showed no change in endoderm muscle activity with osmolarity. CB neurons localize within the ectoderm layer, so their activity and those of ectoderm muscle are mutually consistent (Figure 2 and 3). Thus, CB neurons could be the motor neurons that forms synapse onto ectodermal muscle cells and activate them. On the other hands, endoderm muscle appears not to contact CB neurons or ectoderm muscle (Rushforth and Burke 1971, Dupre and Yuste 2017), behaving as a separate system, unaffected by osmolarity. Future experiments could examine ectoderm and endoderm muscle activity together, with simultaneous calcium imaging of both tissues with two different color indicators. Also, simultaneous imaging of neurons and muscle cells using transgenic *Hydra* that expresses different color calcium sensors in both sets of cells, could explore the relationship between CB neurons and ectoderm muscle.

**Figure 4.**
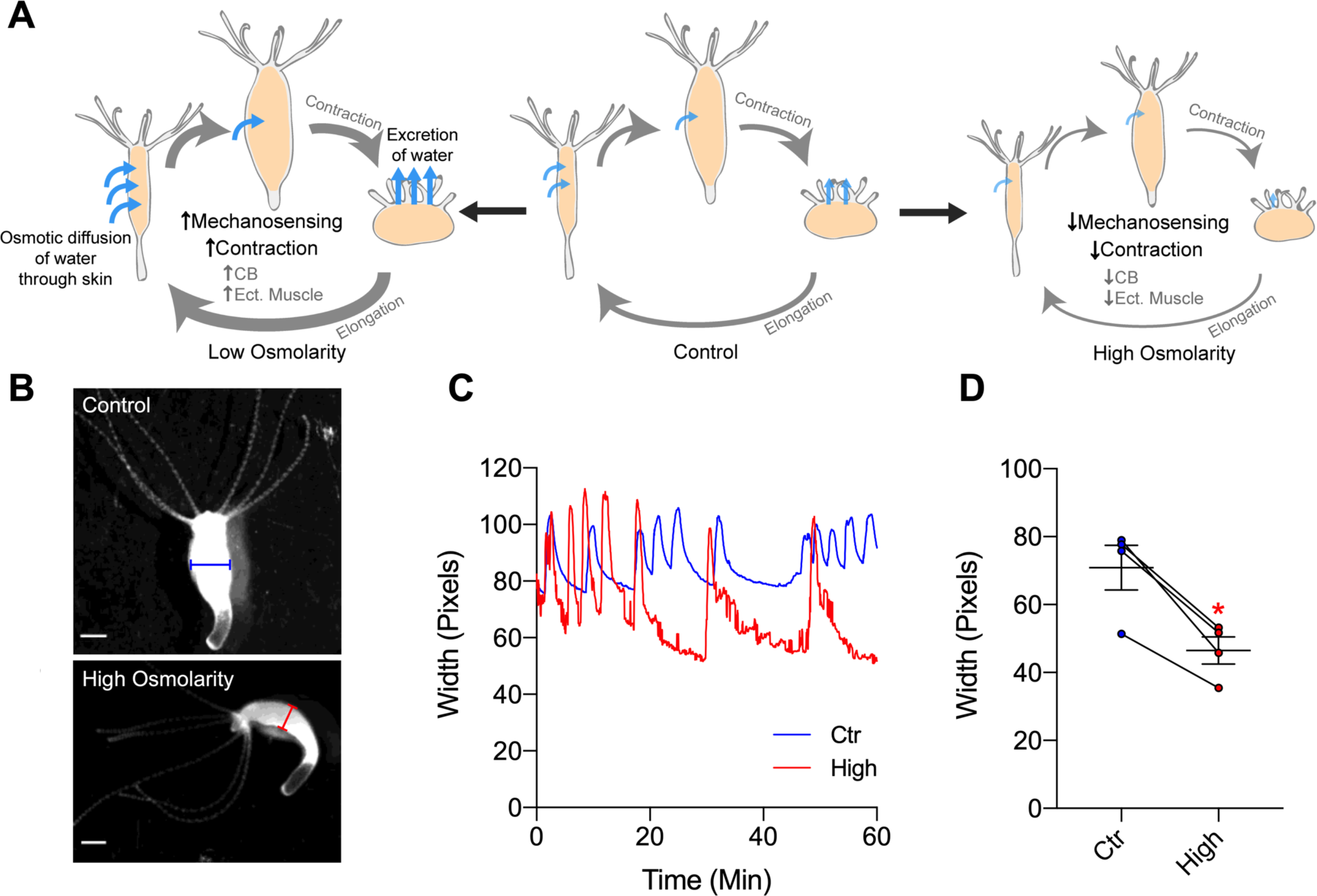
Model explaining *Hydra’s* response to osmolarity and effect on body width. ***A***, Schematic model depicting how *Hydra* changes body width depending on osmolarity. Light-blue arrows indicate the direction and speed of water accumulation, which swells *Hydra*’s body and activate mechanosensory system and contractions. ***B***, Representative images show width of *Hydra*’s body column at the end of elongation cycle, under control media (blue, above) or high osmolarity solution (red, below). ***C***, Representative traces show changes in width over time under control media (blue) or high osmolarity solution (red). ***D***, Width of body column in control media (blue, 70.962 ± 6.560) or high osmolarity solution (red, 46.540 ± 4.036). Line depicts the same animal in each condition. Error bars are shown as the mean ± SEM, with symbol marks denoting data points from individual *Hydra* (*N* = 4). Student t test was performed: \**p* < 0.05.

We found conditions that changed contractions in free behavior without altering neuronal or muscle activity in mounted preparations. For instance, during free behavior, high temperature (30°C) increased the number of contractions and foot detachments (Fig. 1E and F). Above 25°C, *Hydra* activates heat shock protein pathways leading to apoptosis. 30°C is eventually lethal to *Hydra* (Bosch, Krylow et al. 1988), so increased locomotion could reflect an escape behavior, one absent in mounted preparations. We also found that well-fed animal (4 shrimp per day), had fewer contractions, but increased foot detachments (Fig. 1E). It is not clear what would be the function of these behaviors and why these conditions did not alter the activity of neurons or muscles in mounted preparations. Mounting *Hydra* for calcium imaging may have disrupted physiological responses of neurons and muscles in the heat and food. This effect should be reexamined by imaging neurons and muscle activity of freely moving *Hydra*, perhaps with wide-field 3D high-speed scanning systems (Cong, Wang et al. 2017, Kim, Kim et al. 2017).

In summary, using *Hydra*, we developed methods to measure and analyze the activity of the entire neuronal and muscle tissue in an animal during behavior. We find that osmolarity controls the activity of a selective group of neurons and muscle cells, without affecting others, leading to changes in contractile behavior. This approach, measuring the entire neuronal and muscle activity during a simple behavior in an accessible preparation, could be used systematically in *Hydra* and other animals to understand how neuronal and muscle function generates behavior.

## Acknowledgements

We thank H. Shuting for the MATLAB codes and other members of the Yuste Lab and the MBL *Hydra* Lab for assistance and A. Fairhall for discussions.

## Authors Contributions

W.Y. designed the study and collected and analyzed data. W.Y. and R.Y. interpreted data and wrote the manuscript. R.Y. directed the study and secured equipment and funding.

## Supplemental movies

**Movie 1. Freely-moving *Hydra* in control media.** Animals were allowed to move freely in a petri dish. Video was taken at 2 Hz, and sped up 40 fold. Scale bar, 1 mm.

**Movie 2. Ectoderm muscle activity in control media.** The animal was allowed to move between coverslips in mounted configuration. Video was taken at 2 Hz, and sped up 20 fold. Scale bar, 500 µm.

**Movie 3. Neural activity in control media.** The animal was allowed to move between coverslips in mounted configuration. Video was taken at 2 Hz, and sped up 20 fold. Scale bar, 500 µm.

